# An engineered tumor organoid model reveals cellular identity and signaling trajectories underlying translocation RCC

**DOI:** 10.1101/2023.09.01.554626

**Authors:** Maroussia M.P. Ganpat, Francisco Morales-Rodriguez, Nhung Pham, Philip Lijnzaad, Terezinha de Souza, Sepide Derakshan, Arianna Fumagalli, Peter Zeller, Aleksandra Balwierz, Dilara Ayyildiz, Marry M. van den Heuvel-Eibrink, Ronald R. de Krijger, Alexander van Oudenaarden, Thanasis Margaritis, Susana M. Chuva de Sousa Lopes, Jarno Drost

## Abstract

Translocation renal cell carcinoma (tRCC) is a rare, aggressive type of kidney cancer primarily occurring in children. They are genetically defined by translocations involving MiT/TFE gene family members, TFE3 or, in rare cases, TFEB. The biology underlying tRCC development remains poorly understood, partly due to the lack of representative experimental models. Here, we utilized human kidney organoids, or tubuloids, to engineer a tRCC model by expression of one of the most common MiT/TFE fusions, SFPQ-TFE3. Fusion expressing tubuloids adopt a tRCC-like phenotype and gene expression signature *in vitro* and grow as clear cell RCC upon xenotransplantation in mice. Genome-wide binding analysis reveals that SFPQ-TFE3 reprograms gene expression signatures by aberrant, gain-of-function genome-wide DNA binding. Combining these analyses with single-cell mRNA readouts reveals an epithelium-to-mesenchymal differentiation trajectory underlying tRCC transformation, potentially caused by deregulated Wnt signaling. Our study demonstrates that SFPQ-TFE3 expression is sufficient to transform kidney epithelial cells into tRCC and defines the trajectories underlying malignant transformation, thereby facilitating the development of new therapeutic interventions.

## Introduction

Translocation renal cell carcinoma (tRCC) is an aggressive subtype of RCC. While in adults tRCC only encompasses about 5% of all RCC cases, it is a particularly common RCC subtype in children comprising up to 50% of cases (1-3). Despite multimodal treatment strategies, outcomes remain dismal for patients with refractory or metastatic disease emphasizing the need to increase our knowledge on tRCC development to facilitate therapeutic innovation. Genetically, tRCCs are uniquely defined by gene fusions involving Melanocyte Inducing Transcription/Transcription Factor E (MiT/TFE) gene family members TFE3 or TFEB. These fusions make tRCC fundamentally different from the majority of RCC, which typically harbor a high mutational load and numerous copy number alterations. Using mouse models, tubular epithelial cells were shown to be permissive to tRCC formation upon expression of the oncogenic fusion (4). Large genome sequencing efforts, however, revealed recurrent genetic alterations in tRCC that could serve as tumor drivers. It therefore remains unknown whether MiT/TFE fusion expression is sufficient to drive tRCC formation (5-8).

RCCs include multiple heterogeneous subtypes that can be further subdivided based on their histological appearance (9, 10). These include clear cell RCC (ccRCC), defined by tumor cells with distinct clear cytoplasms caused by intracellular accumulation of lipids and glycogen, and papillary RCC (pRCC) (11, 12). TRCC comprises both histological subtypes, most likely caused by different cellular origins. By comparing single cell transcriptomes and epigenomes of human RCC with different histological subtypes to gene expression signatures of the developing human kidney, ccRCC were suggested to arise from proximal tubular epithelium and pRCC from more distal parts of the nephron (13-15). An engineered *in vitro* model representative of human tRCC allowing for studying tumor initiation and progression has not been developed yet.

We have previously described the establishment of organoid cultures from normal and cancerous human kidney tissues (16, 17). Normal kidney derived organoids are primarily composed of proximal tubule cells and, to a lesser extent, of cells from the collecting duct and loop of Henle. These so-called tubuloids retain the characteristics of functional renal epithelial cells (17). Normal human tissue-derived organoids were previously used to engineer tumor progression models of different types of cancer by introducing (combinations of) different putative driver mutations (18-21). These models were shown to be representative of patient tumors, allowing for the molecular dissection of tumorigenesis.

Here, we use human tubuloid cultures to engineer a tRCC model representative of human disease. Applying a multi-omics approach to these genetically engineered tRCC organoids, our study reveals fusion-induced signaling and differentiation trajectories underlying tumorigenesis.

## Results

### SFPQ-TFE3 expression is sufficient to transform human kidney tubuloids into ccRCC

Adult stem cell-derived human kidney organoids are primarily composed of tubular epithelium (i.e. tubuloids) (17), the presumed cellular origin of RCC (13, 15). We therefore set out to model tRCC by ectopic expression of one of the most common MiT fusions in tRCC, *SFPQ-TFE3*, in two independent human tubuloid models (Tub^Fus^) using lentiviral transduction (Fig. 1A). As controls, tubuloids were transduced with either a luciferase (Tub^Ctrl^), or wildtype *TFE3* (Tub^TFE3^) expression plasmid (Fig. 1B). Quantitative reverse transcription polymerase chain reaction (qRT–PCR) (Fig. 1C) and western blot analyses confirmed the expression of the intended genes (Fig. 1D). Compared to Tub^Ctrl^ and normal human kidney tubuloids, no difference in appearance was observed in Tub^TFE3^ cultures (Supplementary Fig. 1A). Tub^Fus^ cultures, however, consistently contained a mixture of the typical cystic as well as distinct dense, solid structures. Hematoxylin and eosin (H&E) staining further confirmed that Tub^Ctrl^ and Tub^TFE3^ both appeared as normal tubuloids, with cystic and folded structures composed of a single-layered epithelium (17). Remarkably, Tub^Fus^ cultures were enriched for multilayered and solid organoids containing cells with distinctive clear cytoplasms (Fig. 1B,E), similar to clear cell RCC (ccRCC) (Fig. 1E) (16). In contrast to Tub^Ctrl^ and Tub^TFE3^, Tub^Fus^ cultures showed strong nuclear TFE3 expression (Fig. 1E), another hallmark of tRCC (22). Of note, no apparent increase in proliferation rate could be observed in any of the cultures, as determined by Ki67 immunostaining (Supplementary Fig. 1B).

**Fig. 1.**
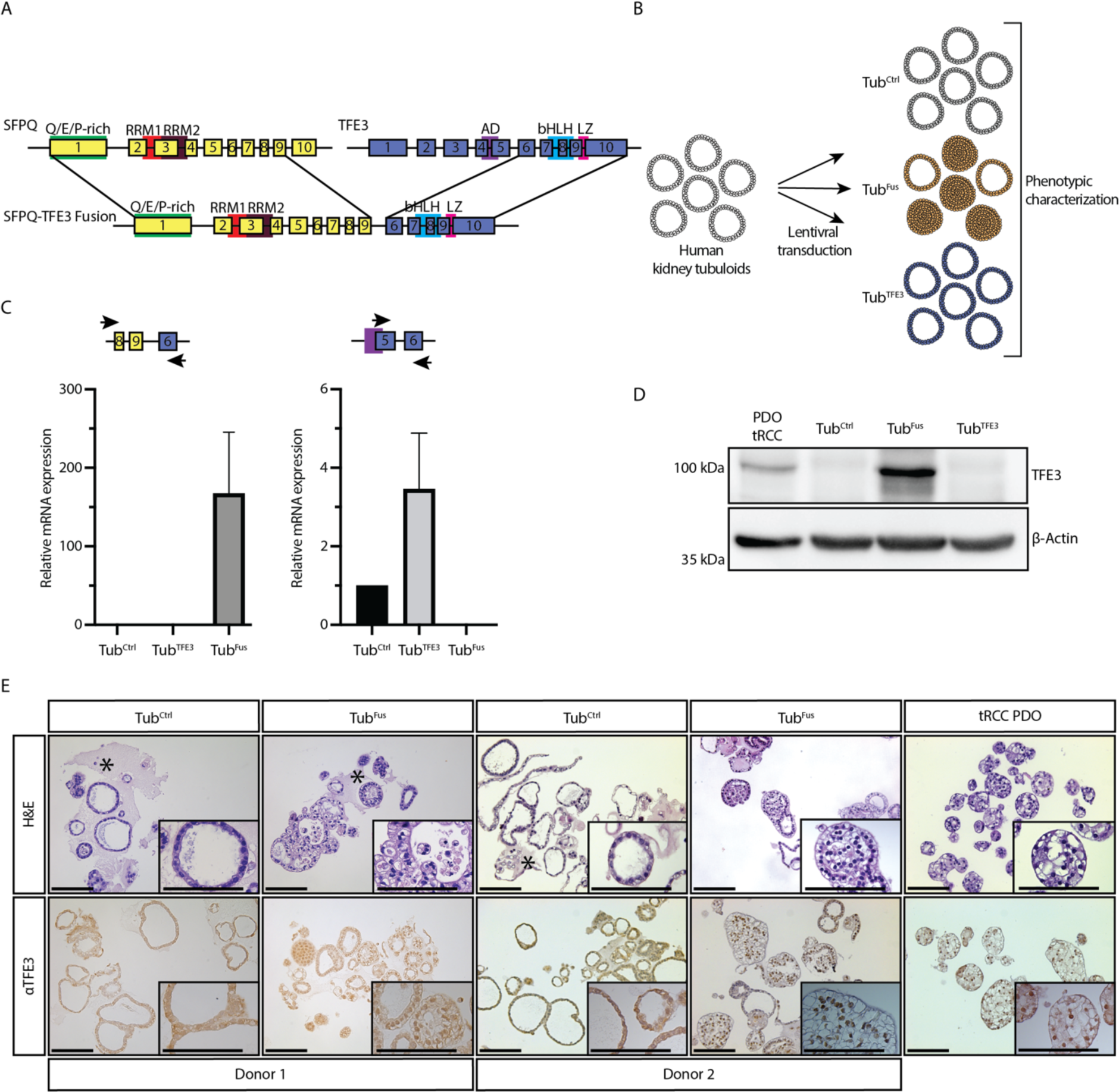
SFPQ-TFE3 fusion induces clear cell RCC phenotype and retains nuclear TFE3 localization. **(A)** Schematic representation of the generated SFPQ-TFE3 gene fusion. Numbers depicting the exons. **(B)** Schematic representation of the experimental setup. Normal human kidney organoids (i.e, tubuloids) are lentivirally transduced with either a control plasmid (Tub^Ctrl^), SFPQ-TFE3 expressing plasmid (Tub^Fus^), or wild-type TFE3 expressing plasmid (Tub^TFE3^). **(C)** qRT-PCR shows expression of *SFPQ-TFE3* fusion and *TFE3* wildtype on mRNA level. Error bars show standard deviation of 3 independent experiments. **(D)** Western blot analysis of SFPQ-TFE3 fusion expression in human tubuloid cultures lentivirally transduced with either a control, SFPQ-TFE3, or TFE3 expression plasmid. β-actin protein levels are used as loading control. Patient-derived tRCC organoids were included as positive control (16). Molecular weight markers are indicated in kilodalton (kDa). **(E)** Histological characterization of the indicated human tubuloid cultures and patient-derived tRCC organoids by H&E and immunostaining for TFE3. Scale bars equivalent to 100 μm. * indicating residual BME. PDO: Patient-derived organoid. tRCC: translocation renal cell carcinoma. Q: glutamine. E: glutamic acid. P: proline. RRM: RNA-recognition motif. AD: activation domain. bHLH: basic helix loop helix. LZ: leucine zipper. H&E: hematoxylin & Eosin.

Next, we investigated whether our engineered Tub^Fus^ organoids are tumorigenic *in vivo* by orthotopic transplantation of Tub^Ctrl^ and Tub^Fus^ into the kidney capsule of immunodeficient mice (Fig. 2A). After 16 weeks, mice were sacrificed, and the kidneys were analyzed for tumor growth. No mice (0/4) transplanted with Tub^Ctrl^ cells developed substantial nodules (Fig. 2B). Histological analysis revealed small grafts that expanded minimally consisting of tubuloid-like cysts lined by single layered epithelium that stained positive for a human-specific cytokeratin (hKRT), thus confirming its human origin (Fig. 2C). In contrast, all mice (5/5) transplanted with Tub^Fus^ organoids developed tumors displaying histological features of ccRCC (Fig. 2B,C). In addition, TFE3 IHC staining showed apparent nuclear localization in Tub^Fus^-derived tumors. Thus, introduction of the MiT fusion *SFPQ-TFE3* enables normal human kidney tubuloids to grow as tumors with ccRCC features *in vitro* and *in vivo*.

**Fig. 2.**
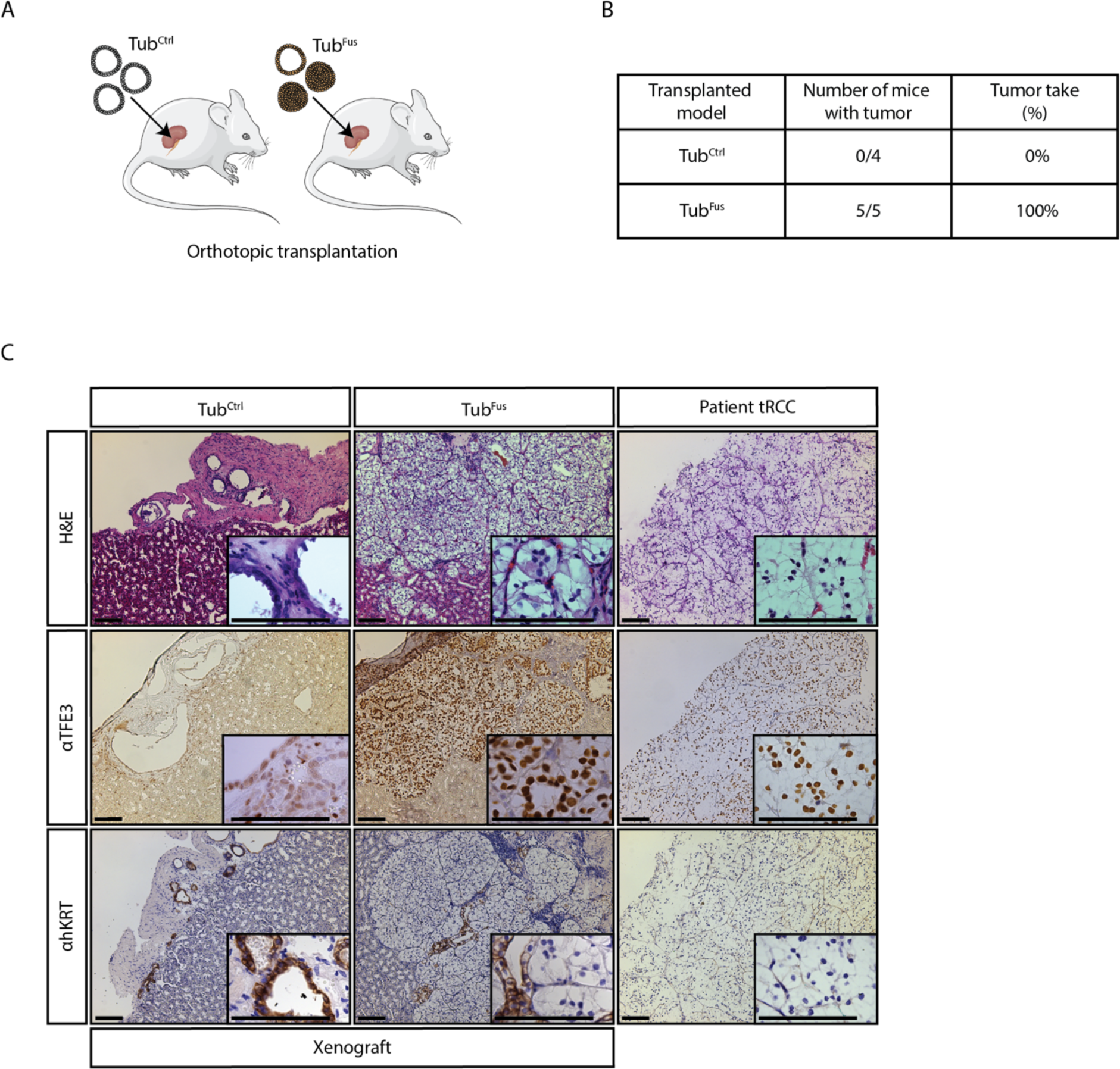
SFPQ-TFE3 expressing tubuloids grow as tumors with ccRCC features upon orthotopic transplantation in mice. **(A)** Schematic overview of the experiment. **(B)** Overview of tumor development in xenotransplanted mouse models. **(C)** Histological characterization of xenograft-derived and tRCC patient-derived tissue by H&E and immunostaining for TFE3 and hKRT. Scale bars equivalent to 100 μm. H&E: Hematoxylin & Eosin. tRCC: translocation renal cell carcinoma. hKRT: human keratin.

### Tub^Fus^ cells adopt an RCC-like gene expression signature caused by aberrant genome wide binding of SFPQ-TFE3

Having generated a representative model of human tRCC and their patient-matched normal counterparts, we set out to study the oncogenic effects of SFPQ-TFE3. First, we performed bulk mRNA-sequencing (RNAseq) on Tub^Ctrl^, Tub^TFE3^ and Tub^Fus^ cultures, as well as Tub^Fus^-derived tumor xenografts (Fig. 3A). We subsequently compared the gene expression profiles to a previously generated RNAseq dataset of pediatric kidney cancer organoids, which includes several tRCC (16). Unsupervised hierarchical clustering showed that, in line with their phenotypic appearance, Tub^Ctrl^ and Tub^TFE3^ both clustered with normal kidney tubuloid cultures (Fig. 3A). Of note, although increased TFE3 mRNA expression could be detected in Tub^TFE3^ cells (Fig. 1C), no significant changes in gene expression profile were observed when compared to Tub^Ctrl^ cultures (Fig. 3A). Strikingly, Tub^Fus^ organoids and Tub^Fus^-derived xenografts clustered with tRCC organoids, corroborating their histological resemblance to tRCC (Fig. 3A).

**Fig. 3.**
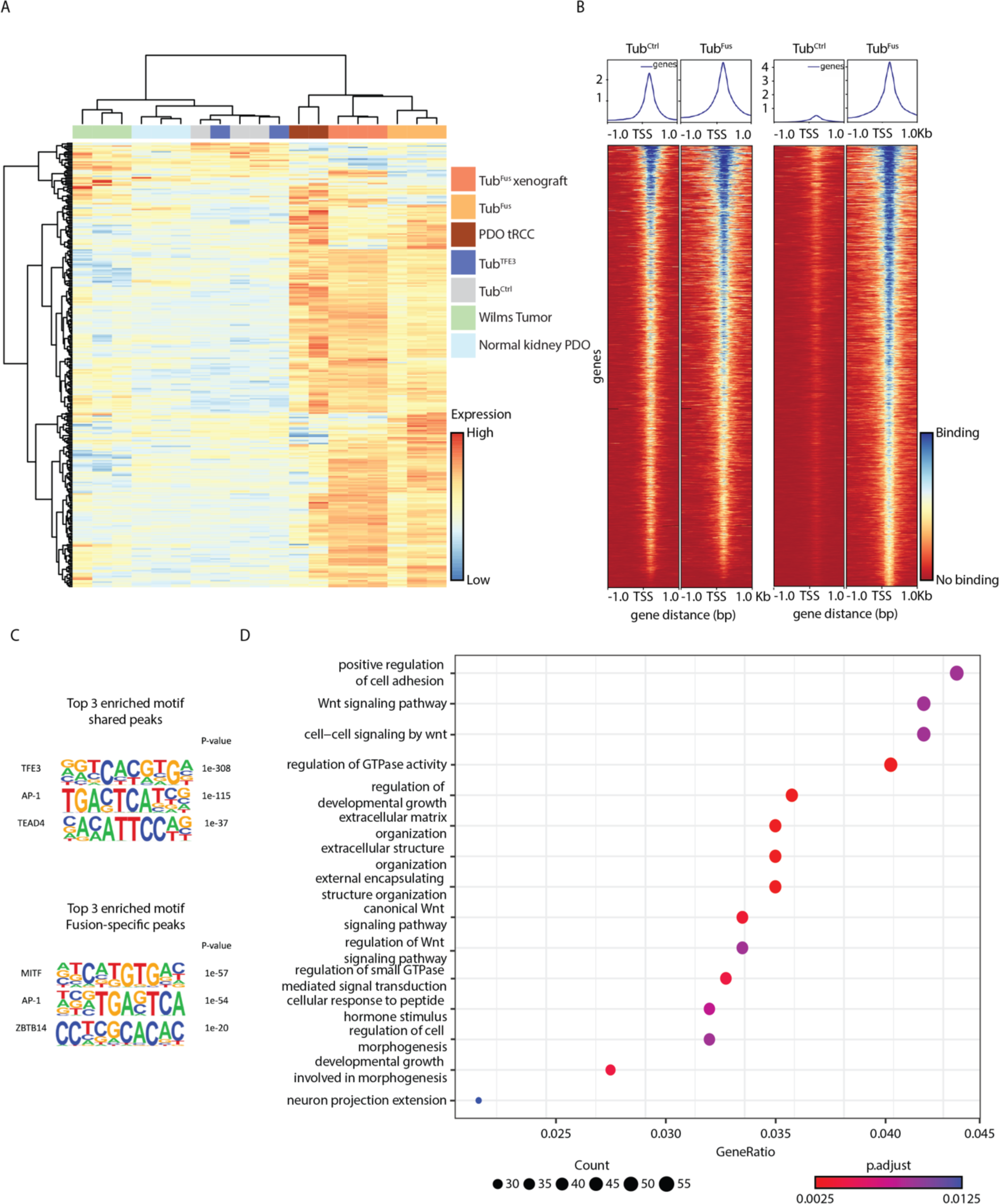
Fusion expressing tubuloids adopt an RCC-like gene expression signature caused by aberrant genome wide binding of SFPQ-TFE3. **(A)** Unsupervised hierarchical clustering of normalized counts from differentially expressed genes detected between Tub^Ctrl^ and Tub^Fus^. Counts were scaled per row. **(B)** Tornado-plots showing the shared peaks and the Tub^Fus^ specific peaks. **(C)** Enriched top-3 motifs of shared peaks and top-3 enriched motifs which are Tub^Fus^-specific. **(D)** Gene set enrichment analysis of fusion-specific peaks.

Previously, chimeric transcription factors were demonstrated to exert their oncogenic effects through aberrant binding to the DNA, thereby dramatically reshaping the gene expression landscape (23, 24). To investigate which genes are directly regulated by the SFPQ-TFE3 fusion, we set out to map genome-wide binding of wildtype TFE3 and the SFPQ-TFE3 fusion using CUT&RUN (25). Our models give us the unique opportunity to do so, as we can compare the engineered cancer cells (Tub^Fus^) to their matching normal counterparts (Tub^Ctrl^). We found 2,774 peaks shared between Tub^Ctrl^ and Tub^Fus^, and 2,653 peaks to be specific for Tub^Fus^ (Fig. 3B, Supplementary Fig. 2A, Supplementary Table 1,2). Notably, we did not find any Tub^Ctrl^-specific TFE3 peaks, suggesting that at least the vast majority of detected shared peaks are bound by endogenous wildtype TFE3. Peak annotation showed that there is no apparent change in genome-wide distribution between peaks in Tub^Ctrl^ and Tub^Fus^ (Supplementary Fig. 2B). *De novo* motif analysis revealed that the peaks shared between Tub^Ctrl^ and Tub^Fus^ were, as expected, enriched for the TFE3 E-box CLEAR (coordinated lysosomal expression and regulation) element (Fig. 3C). Interestingly, Tub^Fus^-specific peaks were highly enriched for the MITF motif (Fig. 3C), suggesting that the oncogenic fusion has gain-of-function activities by binding to genomic regions beyond the typical TFE3 binding sites. Gene set enrichment analysis (GSEA) on Tub^Fus^-specific enriched peaks revealed enrichment of genes connected to the Wnt signaling pathway (Fig. 3D). Of the genes bound by the fusion, we find 75 genes to be differentially expressed in Tub^Fus^ compared to Tub^Ctrl^ (24.3% of differentially expressed genes) (Supplementary Fig. 2C, Supplementary Table 3), suggesting that these genes are direct transcriptional targets of the fusion. Thus, fusion expression induces a gene expression profile similar to tRCC, which is at least partly caused by direct binding of the fusion to the target genes and subsequent regulation of their transcription.

### Single cell transcriptomics reveals fusion-driven trajectories underlying malignant transformation

Upon SFPQ-TFE3 expression, we find a mixture of transformed and non-transformed tubuloids (Fig. 1E), suggesting that a certain population of cells is permissive to malignant transformation upon SFPQ-TFE3 expression. To identify this population and define the signaling networks contributing to their transformation, we subjected Tub^Ctrl^ and Tub^Fus^ organoids to single-cell mRNA-sequencing (scRNAseq) using the Chromium 10x Genomics platform. After filtering, 863 cells were left for further analyses (Supplementary Fig. 3A). Projecting cells in t-distributed stochastic neighbor embedding (t-SNE) plots revealed that Tub^Ctrl^ and Tub^Fus^ are mostly similar (Fig 4A). Comparing single cell transcriptomes of the transduced organoids to single cell transcriptomes of the developing kidney (26), we found that Tub^Ctrl^ are primarily composed of tubular epithelium arising from different nephron segments (Fig. 4B), as described before (17). Remarkably, cells exhibiting signals of proximal connecting tubule (CNT)/principal cells/proximal epithelium were highly underrepresented in Tub^Fus^ (Fig. 4C), suggesting that this (or their direct progeny) is the population of cells that change to a tRCC-like gene expression profile upon fusion expression. This is in line with previous reports pointing towards proximal tubular epithelium as the cellular origin of ccRCC (15). Instead, we found a distinct cluster of cells highly enriched in Tub^Fus^ with signals similar to stromal progenitor cells (Fig. 4B,C). These cells were marked by expression of known markers of tRCC, such as *GPNMB* (Supplementary Fig. 3C) (4) and *CTSK* (Supplementary Fig. 3D) (27), strongly suggesting that this the population that transformed to tRCC.

**Fig. 4.**
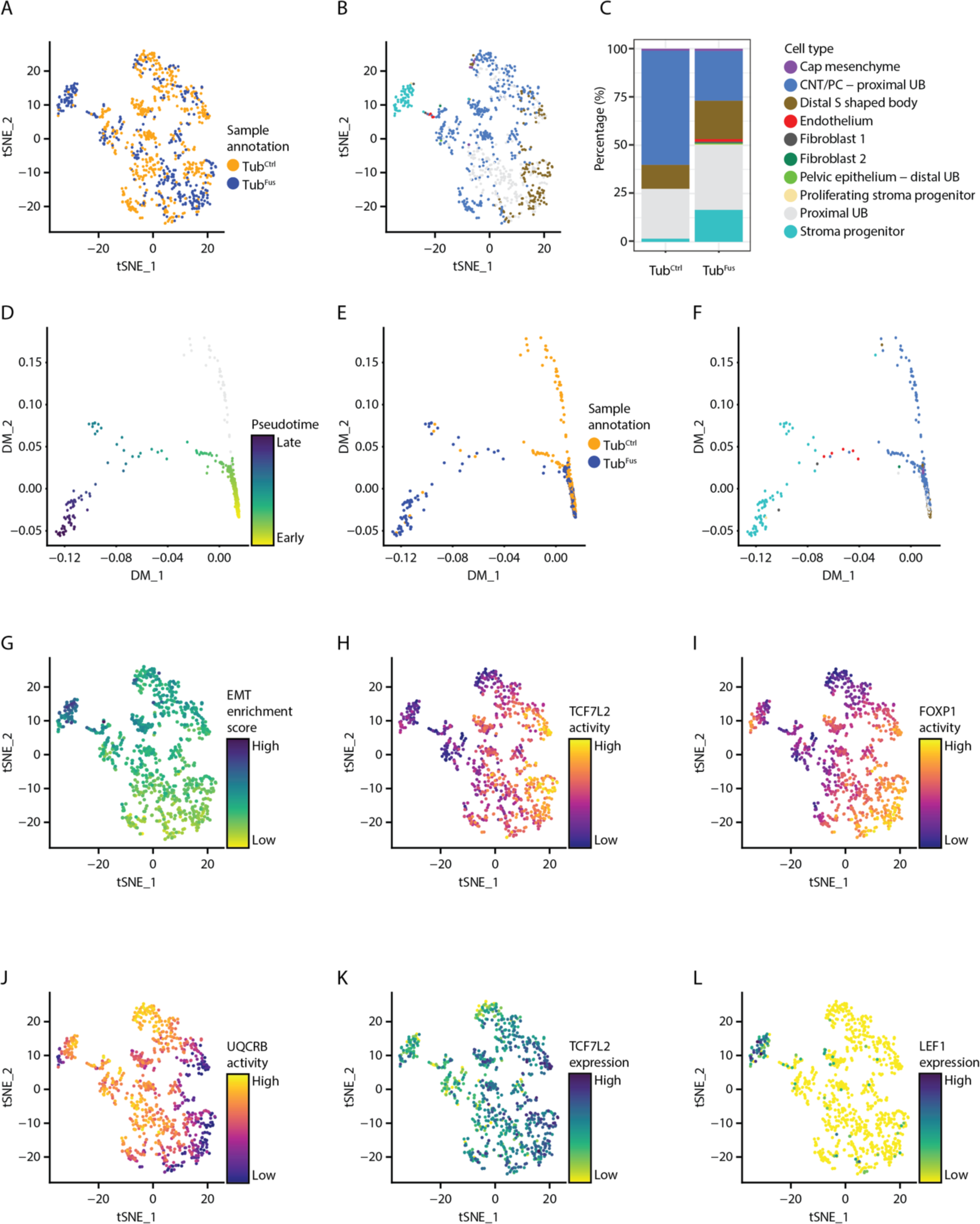
scRNAseq analysis shows fusion-induced trajectories and stromal differentiation of Tub^Fus^ tubuloids. **(A)** t-SNE projection of transduced tubuloids, colored by cluster assignment. **(B)** t-SNE projection of cell typing, colored by cell type assignment. **(C**) Bar graph showing the distribution of the different cell types found in Tub^Ctrl^ and Tub^Fus^, as well as the distribution of the different cell types in the nephron and stromal population. **(D)** Diffusion map showing pseudotime projection, colored by pseudotime. **(E)** Diffusion maps projecting sample annotation, colored by cell cluster. **(F)** Diffusion map projection of transduced tubuloids, colored by cell type. **(G)** t-SNE projection of EMT, colored by enrichment score. **(H)** t-SNE projection of *UQCRB* transcription factor activity, colored by activity levels. **(I)** t-SNE projection of *FOXP1* transcription factor activity, colored by activity levels. **(J)** t-SNE projection of *TCF7L2* transcription factor activity, colored by activity levels. **(K)** t-SNE projection of *TCF7L2* expression, colored by expression level. **(L)** t-SNE projection of *LEF1* expression, colored by expression level.

Next, we investigated the cell state trajectories underlying fusion-induced malignant transformation. To do so, single cell transcriptomes were projected in diffusion space and directionality assessed using pseudotime (Fig. 4D-F). This resulted into two trajectories of which one, referred to as pseudotime 1, corresponds to the differentiation trajectory of tubular epithelium towards the stromal-like, presumed tRCC population (Fig. 4D,F). The tRCC trajectory was further confirmed by increased expression of *GPNMB* along pseudotime 1 being highly enriched in the stromal-like cells compared to the nephron-like progenitor population (Supplementary Fig. 3B,C,E). Mapping cellular processes by GSEA revealed epithelial-to-mesenchymal transition (EMT) to be significantly upregulated in this tRCC population (Fig. 4G, Supplementary Fig. 4A), which is exemplified by high expression of collagens such as *COL3A1, COL6A1 and COL6A3* (Supplementary Fig. 4B), and downregulation of epithelial markers such as keratins (*KRT8, KRT18* and *KRT19)* (Supplementary Fig. 4C). Using single-cell regulatory network interference and clustering (SCENIC), we found increased transcriptional activity of the reactive oxygen species (ROS) factor *UQCRB* (Fig. 4H), suggesting a potential role for ROS in the stromal cells. Interestingly, we found decreased transcriptional activity of the transcription factor *FOXP1* (Fig. 4I) and the Wnt pathway transcription factor *TCF7L2* (Fig. 4J), the latter being in line with our CUT&RUN analyses (Fig. 3D, Supplementary Fig. 2D). This coincided with decreased *TCF7L2* mRNA expression in the stromal population (Fig. 4K). Interestingly, expression of *LEF1* (Fig. 4L), another key Wnt pathway transcription factor, is markedly upregulated in the stromal population, suggesting that a switch in TCF7L2/LEF1 transcription factor binding is contributing to the oncogenic properties of the SFPQ-TFE3 fusion in tRCC. Such a switch was previously shown to be critical for differentiation of nephron progenitor cells (28). Altogether, we generated an organoid model for tRCC that is representative of human disease revealing differentiation trajectories and underlying signaling pathways that could be therapeutically explored.

## Discussion

We have generated the first engineered human tRCC organoid model. This enabled us to demonstrate that expression of the MiT/TFE fusion *SFPQ*-*TFE3* is sufficient to transform normal human renal tubular epithelial cells to tRCC. Our data provides the first functional evidence in a human setting that cells coming from the proximal part of the nephron (i.e., proximal connecting tubule, principal cells, and proximal epithelium) are the cells being permissive to malignant transformation upon fusion expression. This is in line with previous reports based on comparing tumor transcriptomes with normal kidney expression profiles (13, 15) as well as data from transgenic mouse models using a tubular epithelium specific Cre driver of fusion expression (4).

Analyzing single cell transcriptomes of fusion-expressing tubuloids, we found that the presumed tRCC population (expressing biomarkers of tRCC) is highly enriched for stromal markers and depleted from epithelial markers indicative of an EMT. This phenomenon has also been observed in patient RCC, as tumors typically display decreased expression of epithelial markers such as keratins and increased expression of collagens (29). Notably, during nephrogenesis, mesenchymal to epithelial transition (MET) is responsible for the development of functional nephrons from nephron progenitor cells residing in the cap mesenchyme. The Wnt signaling pathway is critical for this transition from mesenchyme to epithelium (30, 31). Previously, kidney-specific overexpression of wild-type *TFEB* in transgenic mice was shown to induce ccRCC and pRCC, at least partially, through enhanced Wnt signaling (32). Intriguingly, we found downregulation of Wnt pathway activity along the trajectory of transitioning from tubular epithelium to the tRCC-like stromal cells. These different observations could be explained by the activity of different MiT transcription factors (wild-type *TFEB* versus oncogenic chimeric *SFPQ*-*TFE3*) and therefore different tRCC subtypes, or by the differences between mouse and human disease models. We also find decreased expression of *RNF43*, a negative regulator of Wnt signaling (33), upon fusion expression, which was previously shown to be an indicator of poor prognosis of RCC patients (34), suggesting that our model represents the more aggressive tRCC subtype.

We show that the oncogenic SFPQ-TFE3 fusion regulates gene transcription beyond the canonical TFE3 targets by gain-of-function binding to the MITF consensus motif, very similar to the TFE3 E-box CLEAR consensus motif. Similar observations were made for other oncogenic chimeric transcription factors such as *EWS-FLI1* in Ewing sarcoma (24, 35). By doing so, oncogenic chimeric transcription factors can activate transcription of tumor-specific neotranscripts that are translated to neopeptides or neoproteins, presented on the cell surface of tumor cells, and, as such, recognized by the immune system (24). Immune checkpoint inhibitors were demonstrated to be effective in a subset of tRCC, despite their low mutation burden (36). A potential role for MiT fusion-induced neoantigen expression should be further explored. Our engineered tRCC model can be instrumental in the identification of such transcripts, as it allows for comparing the tumor transcriptome to the transcriptome of its genetically identical normal cell counterpart.

## Acknowledgments

We thank the clinical teams and anonymous volunteer tissue donors that facilitated our research, the Princess Máxima Center Single Cell Genomics facility and Oncode’s Single-cell (epi)-genome sequencing facility (location Hubrecht Institute) for supporting scRNAseq and CUT&RUN, the Princess Máxima Center diagnostic department and Harry Begthel (Hubrecht Institute) for immunohistochemical analyses.

## Materials and Methods

### Ethical statement

The Medical Ethical Committee of the Leiden University Medical Center granted approval of this study (P08.087). Based on the Declaration of Helsinki (World Medical Association), informed consent was obtained.

### Tissue processing and establishment of human kidney tubuloid cultures

Elective abortion material (without medical indication) containing human fetal kidneys at gestational age week 16 (named Donor 1) and 17 (named Donor 2) were collected. Following collection, the samples were cleaned with 0.9% NaCl (Fresenius Kabi, France) and kept on ice until it was processed further (37).

Collected fetal kidneys were minced and digested in AdvDF+++ (Advanced DMEM/F12 containing 1× Glutamax, 10 mM HEPES and antibiotics) containing 1 mg/ml collagenase (Sigma, C9407) and 10 µM Y-27632 on an orbital shaker for 30 - 45 min at 37 °C. The suspension was subsequently washed with AdvDF+++ followed by centrifugation at 250×g. The cell pellets were seeded in growth factor-reduced BME (Trevigen, 3533-010-02) and cultured in kidney organoid medium (AdvDF+++ supplemented with 1.5% B27 supplement (Gibco), 10% R-spondin-conditioned medium, EGF (50 ng/ ml, Peprotech), FGF-10 (100 ng/ml, Peprotech), N-acetylcysteine (1.25 mM, Sigma-Aldrich), Rho-kinase inhibitor Y-27632 (10 µM, Abmole), A83-01 (5 µM, Tocris Bioscience) and Next Generation Surrogate (NGS) Wnt (0.5 nM, ImmunoPrecise Antibodies NASDAQ), as previously described (23, 24).

### Lentiviral transductions

Lentiviral transductions were performed as previously described (38) with lentiviruses encoding either pLKO.1-UbC-luciferase-blast (Tub^Ctrl^), pLKO.1-UbC-TFE3-blast (Tub^TFE3^), or pLKO.1-UbC-SFPQ-TFE3-blast (Tub^Fus^) as previously described (38). Two days after transduction, 5 µg/ml blasticidin was added to the culture medium to select for successfully transduced cells.

### Immunohistochemistry

Tissues and organoids were fixed in 4% paraformaldehyde, dehydrated, and embedded in paraffin. Immunohistochemistry was performed according to standard protocols on 4 µm sections. Sections were subjected to H&E and immunohistochemical staining. The following primary antibodies were used for immunohistochemical staining: TFE3 (anti-TFE3 ab93808 (Abcam, 1:1000)), hKRT (anti-cytokeratin clone Cam5.2 (BD Biosciences, 1:100)).

### RNA isolation, cDNA preparation and qRT-PCR

Total RNA was extracted using Trizol reagent (Invitrogen) or using the Macherey-Nagel RNA isolation kit according to manufacturer’s instructions. Extracted total RNA was used to produce cDNA using the GoScript reverse transcriptase system (Promega) and qRT-PCR was subsequently performed using IQ SYBR green mix (Bio-Rad) according to the manufacturer’s instructions. Results were analyzed using the ΔΔCt method and normalized to *GAPDH* housekeeping gene expression. qRT-PCR primer sequences: *SFPQ-TFE3*_Fwd 5‘-GGCGGAGGAGCAATGAAC-3’ *SPFQ-TFE3*_Rev 5‘-CAAGCAGATTCCCTGACACA-3’ *TFE3*-WT_Fwd 5‘-GATGAGATCATCAGCCTGGAGT-3‘ and *TFE3*-WT_Rev 5‘-TCGGTCTCAGAGATCTCCCG-3’ *GAPDH*_Fwd 5‘-TGCACCACCAACTGCTTAGC-3‘ and *GAPDH*_Rev 5‘-GGCATGGACTGTGGTCATGAG-3’.

### Bulk RNA sequencing

Trizol (Invitrogen) was used to extract total RNA in accordance with the manufacturer’s instructions. Bioanalyzer2100 RNA Nano 6000 chips (Agilent, Cat. 5067-1511) were used to assess RNA quality. NEBNext® Ultra™ RNA Library Prep Kit (New England Biolabs) was used, according to manufacturer’s instructions, to prepare the sequencing libraries. Paired-end sequencing was performed on the Illumina NovaSeq (PE150) by Novogene.

Raw data files that passed quality control had their adapters trimmed with trim galore (v.0.65) and subsequently aligned against human genome hg38 using STAR (v.2.7.2) and read counts quantified with featureCounts (v. 1.6.7) using genome annotation from Gencode v. 37.

Raw count files from each sample were then merged into a single matrix and processed in R 4.2.1 using the R package DESEq2. Prior to any downstream analysis the count matrix was filtered to keep only genes with an average of at least 5 read counts across all samples, normalized, and transformed into log2 scale.

Gene ontology analysis was performed using the R package clusterProfiler (v. 4.6.0) using a 0.05 q-value cutoff; heatmaps were generated with pheatmap.

### Western blot

Western blot was performed as previously described (18). The following antibodies were used: anti-TFE3 (ab93808, Abcam, 1:1000) and anti-GAPDH (ab-9485, Abcam, 1:1000).

### CUT&RUN

CUT&RUN experiments were performed using a modified protocol for low cell numbers as described in (25) using the following antibodies: anti-TFE3 (ab93808, Abcam, 1:2000), anti-H3K4me3 (monoclonal antibody G.532.8, ThermoFisher Scientific, Invitrogen, 1:1000), anti-H3K4me1 (recombinant polyclonal antibody, ChIP-Verified, 71095 ThermoFisher Scientific, Invitrogen, 1:100) and H3K27ac (Acetyl-Histone H3 (Lys27) XP® #8173, Cell Signaling technology, 1:100).

Libraries were sequenced using an Illumina NextSeq2000 (2x100bp). Sequencing quality was checked using FastQC (v0.12.1). Raw FASTQ data files that passed quality control had their adapters and short reads (<20bp) trimmed with Trim Galore (v0.6.10) and subsequently aligned against human genome hg38 with Gencode v. 37 annotation using Bowtie2 (v2.5.1) with the ‘end-to-end’ option. Samtools view (v1.16.1) was used to convert aligned sam files to bam files. Thereafter, Picard Tools (v52.0) was used to sort aligned files using the SortSam function and then to remove sequencing duplicates using the MarkDuplicates function. Summary plots of FastQC, alignment, and filtered data were created using MultiQC (v1.14). S3norm was used to normalize sequencing depths and signal-to-noise ratios using the default setting. Inputs for S3norm were bedgraph files with a fixed bin size of 200bp. They were generated using bamCoverage from deepTools (3.5.1) with an effectiveGenomeSize as 2913022398 for hg38.

Peaks were then called by MACS2 (v2.2.7.1) using bdgpeakcall function with normalized bedgraph files (S3norm NBP files). Peaks called from replicates in the same experimental condition (i.e., luciferase and fusion) were merged using samtools merge (v1.16.1). Bigwig files were then generated using the bamCoverage function from deepTools (v. 3.5.1) for peak visualization in IGV. To identify differential peaks across conditions, aligned data and peak files were loaded into Rstudio (v. 4.2.2) and analyzed with the package DiffBind (v. 3.8.4). Differential analysis was performed by taking into consideration the different conditions in the contrast design. Differential peaks were selected based on an FDR < 0.05 and absolute fold change >2. Heatmaps of differential peaks and shared peaks between experimental conditions were generated with plotHeatmap function from deepTools.

Peak regions were annotated using the ChIPSeeker (v. 1.34.1) package. Enrichment analyses were performed with the ClusterProfiler (v. 4.6.0) package. Finally, motifs were identified using Homer findMotifsGenome.pl with -len 10.

Quality control reports and a detailed version of our used analysis pipeline are available upon request.

### Orthotopic transplantations

All mice experiments were performed in accordance with protocols approved by the Institutional Animal Care and Animal Welfare Body Utrecht and the Royal Netherlands Academy of Arts and Sciences (PMC.65.3067.1901.1, AVD3990020173067). Orthotopic transplantation of transduced tubuloids into the kidney capsule were performed as previously described in (39, 40). Briefly, 10-19 weeks old male NOD.Cg-Prkdscid Il2rgtm1Wjl/SzJ (NSG) mice were transplanted with 400,000 transduced tubuloids each and sacrificed 16 weeks post transplantation. The tumor growth was weekly monitored by abdominal palpation. Kidney, liver, and brain tissue were resected and analyzed for tumor take and/or presence of metastasis.

### Single cell RNA-sequencing

Transduced tubuloids were dissociated into single cell suspensions using TrypLE Express (ThermoFisher Scientific) supplemented with Rho-kinase inhibitor Y-27632 (10 µM, Abmole) and filtered using a 70 μm filter. Single cell suspensions were labeled using Cell Multiplexing Oligos (CMO) (‘cellplex’) for Single Cell RNA Sequencing with Feature Barcode technology, according to manufacturer’s instructions. The viable single cells from the transduced single cell suspension were sorted based on forward/back scatter properties and the combination of DRAQ5 and DAPI staining on a Sony SH800S cell sorter using a 100 μm filter chip. Chromium Next GEM Single Cell 3‘ (10x Genomics) was performed according to manufacturer’s instructions.

The scRNAseq data were processed using “cellranger” (version 7.1.0, 10X Genomics) using the company’s refdata-gex-GRCh38-2020-A transcriptome (41). Due to poor amplification of the cellplex library, additional cellplex reads were recovered using a custom script that is available upon request. Reads originating from Donor 1 were kept as determined by SNV-based genotype multiplexing using “souporcell” (Singularity image created 1 December 2021) (42). Further processing and analyses were done using Seurat (version 4.3.0) using standard settings (43). Barcodes with fewer than 2000 or more than 100,000 transcripts were discarded, as well as barcodes with a mitochondrial percentage >50%. The following genes were excluded from the list of highly variable genes as being potentially confounding: genes specific to S- or G2M phases, genes correlating with S- or G2M phases, male- and female-specific genes, stress-related genes and ribosomal protein genes (44). Cell typing was done by projection of transduced tubuloid cells into the (non-immune) fetal kidney data set, generated by Stewart et al., (26, 43) which was preprocessed using the same steps as used for our tRCC scRNAseq data set. GSEA was done using R package “clusterProfiler” and the MSigDB file “h.all.v2023.1.Hs.symbols.gmt” (45, 46). Genes subjected to enrichment analysis were required to have a False Discovery Rate (FDR) less than 0.05 and be differentially expressed more than 2-fold. For the purpose of intersecting these lists with genes having CUT&RUN peaks, the latter had to be annotated as “protein_coding” in GENCODE v37 and were required to have an FDR less than 0.05. The pseudotime trajectory was generated using the R packages “destiny”, “mclust” and “slingshot”, all using standard settings (47-49). The analysis of transcription factory activity was done using SCENIC (version 0.11.2), using the raw transcript counts as input data, “hs_hgnc_tfs.txt” as the list of transcription factors, “hg38 refseq-r80 10kb_up_and_down_tss.mc9nr.feather” for the motif database and motif annotation from “motifs-v9-nr.hgnc-m0.001-o0.0.tbl” (50). Inferring the gene regulatory network involves an element of stochasticity to sample the solution space sufficiently, therefore the algorithm was run 20 times and only transcription factors that were found in at least 10 of the runs (50%) were analyzed. Full scripts for the complete preprocessing and downstream analysis steps are available upon request.

## Data availability

Raw bulk RNA-sequencing data and count tables of the single-cell RNA sequencing will be deposited in the European genome-phenome archive. Accession numbers are pending.

## Code availability

All data analysis code will be made available upon request.

## Contributions

M.M.P.G and F.M. performed experiments. S.D. provided technical assistance. N.P. P.L., T.d.S., P.Z., D.A., A.v.O. and T.M. analyzed the sequencing data. A.F. performed the orthotopic transplantations. R.R.d.K. performed pathological review of organoids and xenografts. P.Z. and A.O. helped with CUT&RUN experiments and analyses. M.M.v.d.HE. and S.C.d.S.L. shared resources and clinical expertise. J.D. conceived and supervised the study. M.M.P.G and J.D. wrote the manuscript, with input from all authors. All authors read and approved the manuscript.

## Supplementary figure legends

**Supplementary Fig. 1.**
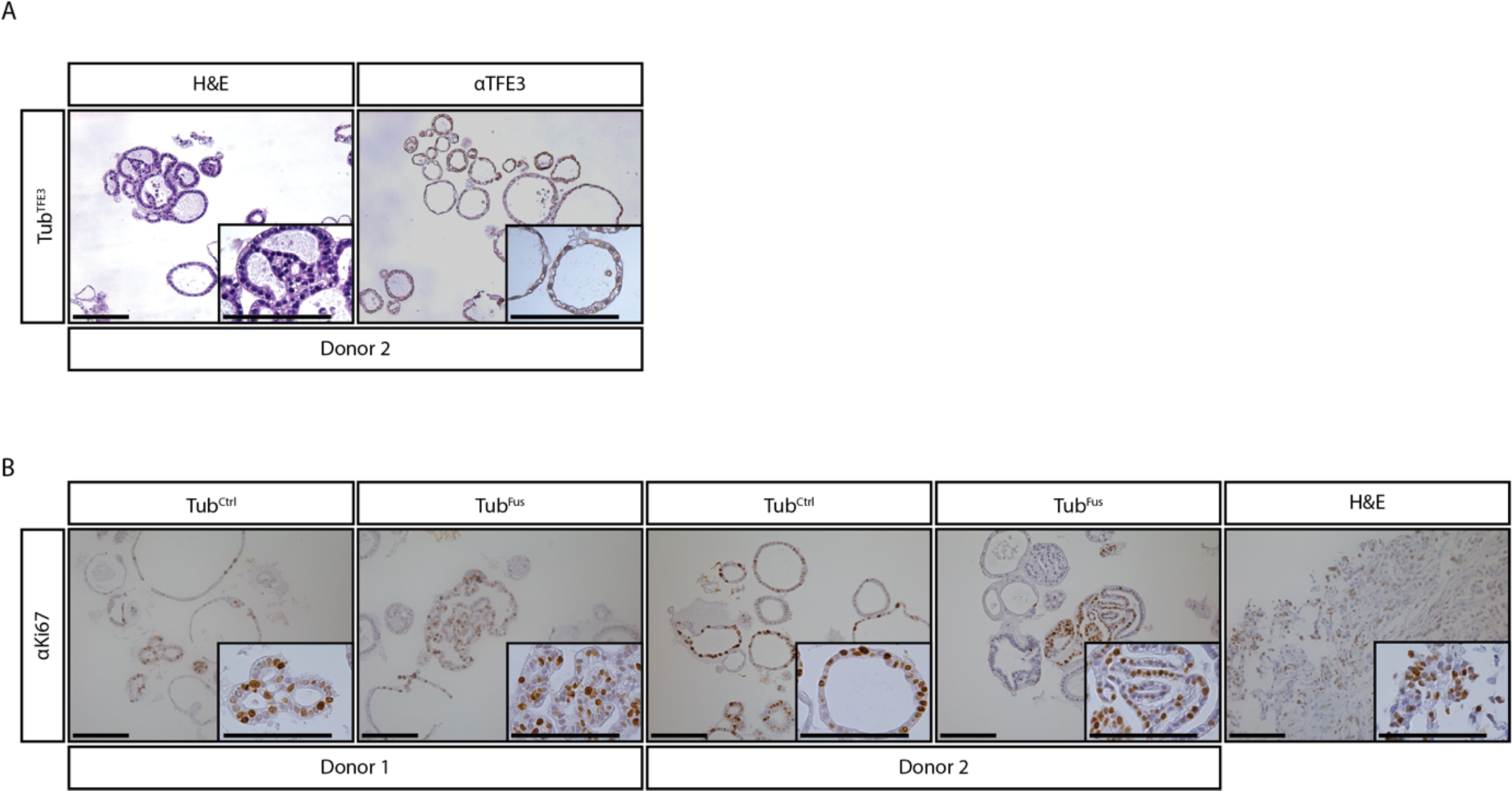
Histological characterization of engineered human tubuloid models. **(A)** Tub^TFE3^ histochemical characterization by H&E and IHC for TFE3. **(B)** IHC staining for Ki67, proliferation assessment. Patient-derived tumor tissue were included as positive control (16). KRT: Keratin. Scale bars equivalent to 100 μm.

**Supplementary Fig. 2.**
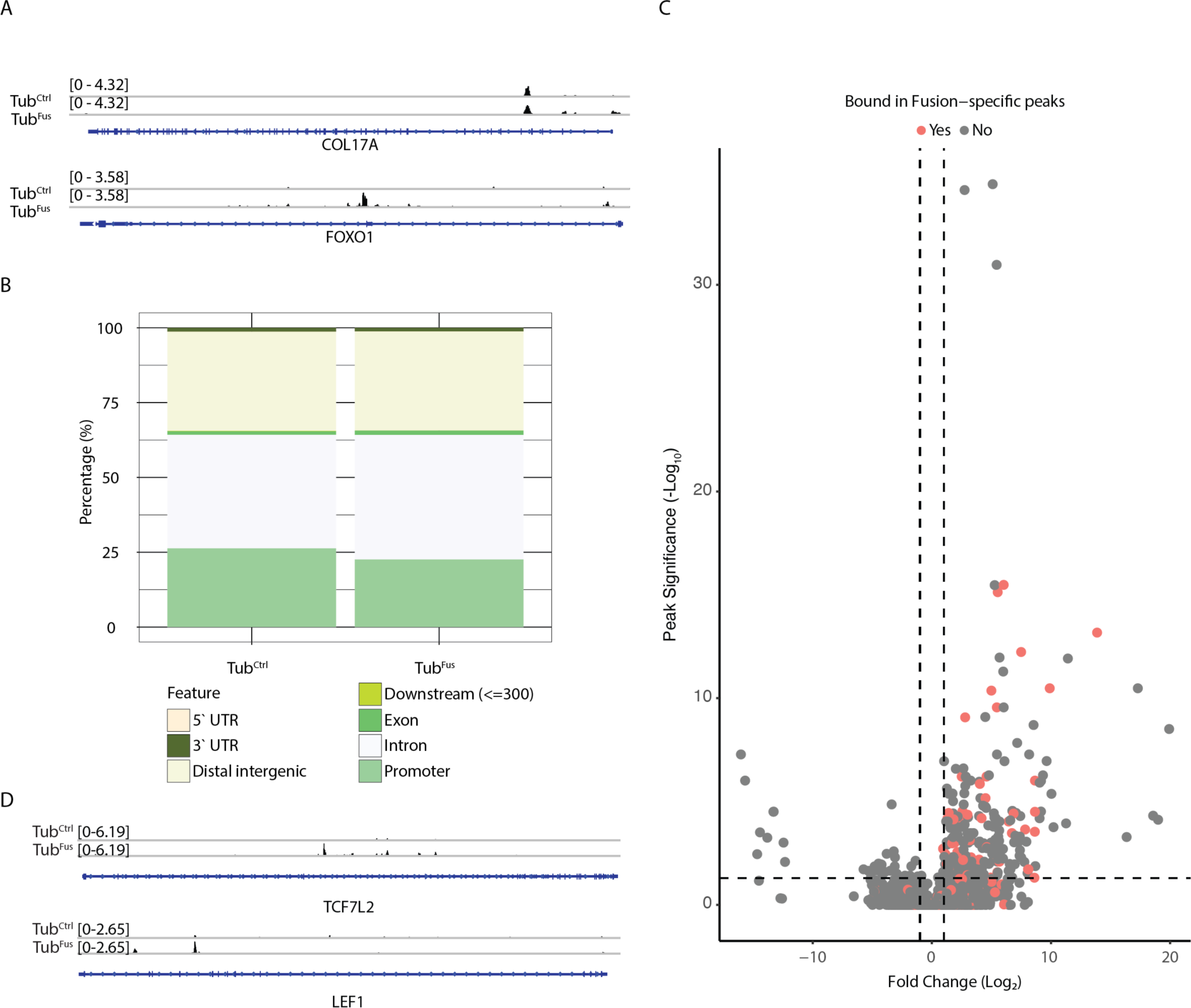
CUT&RUN analyses reveal genome-wide gain of function binding of SFPQ-TFE3. **(A)** Bar graph showing the distribution of the annotated peaks found in Tub^Ctrl^ and Tub^Fus^. **(B)** An example of a shared binding peak between Tub^Ctrl^ and Tub^Fus^ and an example of a fusion-specific binding peak. **(C)** Volcano plot showing the fusion-specific peaks and their corresponding expression on bulk RNAseq. **(D)** Fusion-specific binding of TCF7L2 and LEF1 in Tub^Ctrl^ and Tub^Fus^.

**Supplementary Fig. 3.**
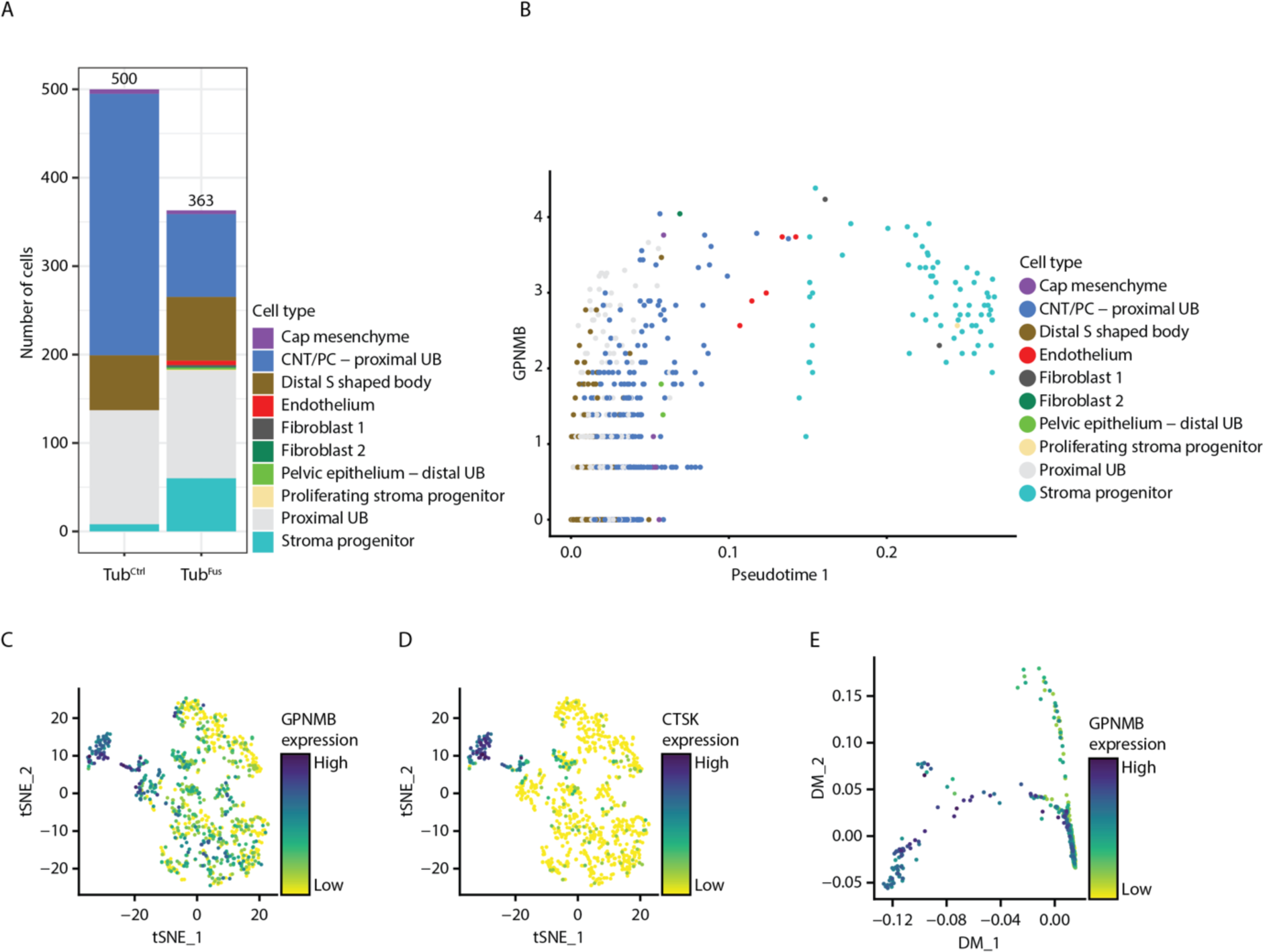
scRNAseq analysis shows fusion-induced trajectories and stromal differentiation in Tub^Fus^ tubuloids. **(A)** Bar graph showing the absolute number of cells per cell type in Tub^Ctrl^ and Tub^Fus^. **(B)** Dot plot showing the expression of *GPNMB* projected onto pseudo trajectory 1. Each dot represents one cell. **(C)** t-SNE projection of the expression of *GPNMB,* colored by expression levels. **(D)** t-SNE projection of the expression of *CTSK*, colored by expression levels. **(E)** Diffusion map projection of *GPNMB* expression along pseudotime 1.

**Supplementary Fig. 4.**
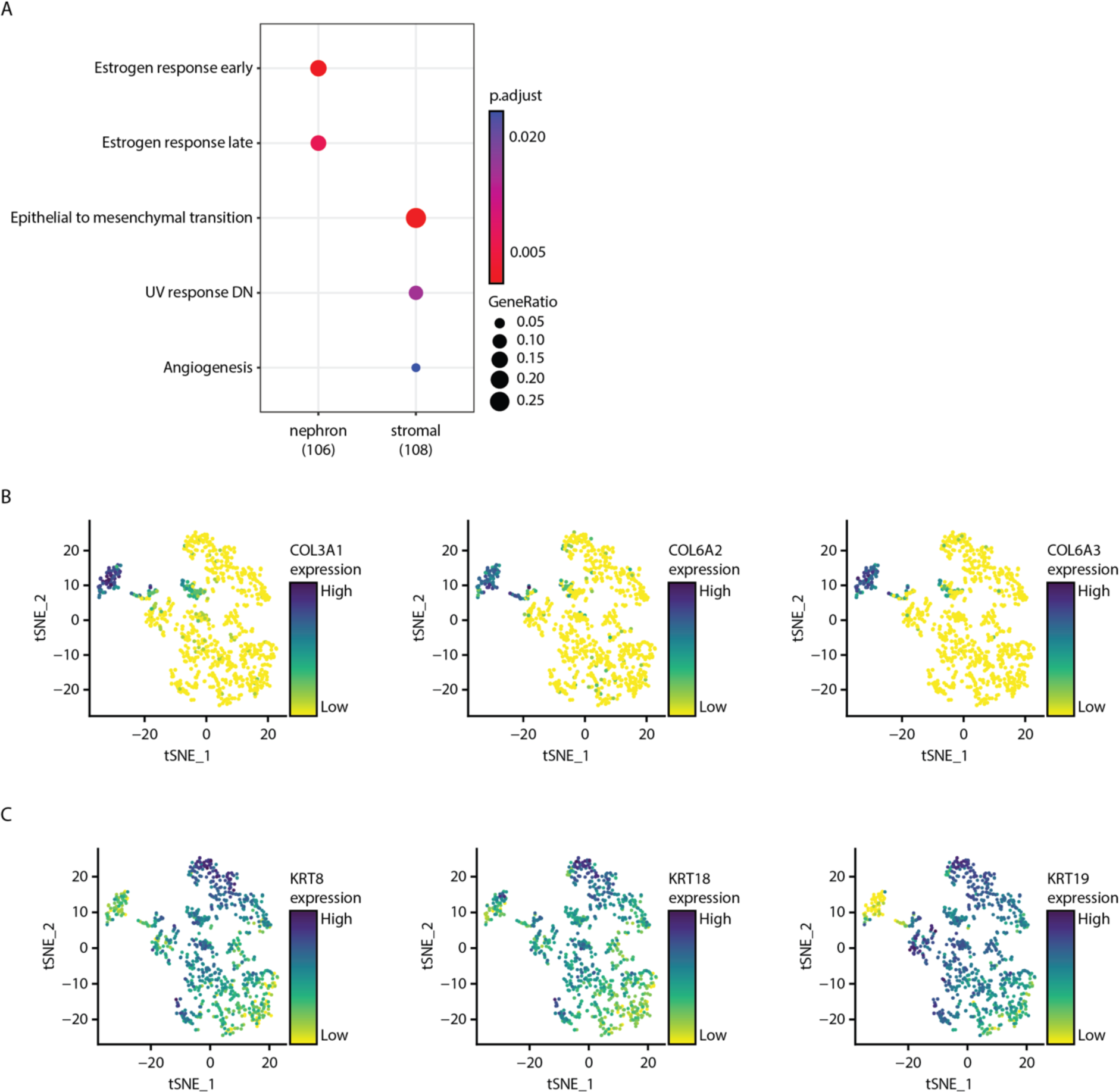
scRNAseq analysis shows involvement of different signaling pathways in the transition to tRCC. **(A)** Enriched biological processes in the nephron and stromal population. **(B)** t-SNE projection of several collagens, i.e., *COL3A1, COL6A2* and *COL6A3,* colored by expression level. **(C)** t-SNE projection of keratins, i.e., *KRT8, KRT18* and *KRT19*, colored by expression level.

## Supplementary table legends

**Supplementary Table 1.** Fusion-specific annotated peaks within 1 and 3 kilobases (kb) of transcription start site (TSS).

**Supplementary Table 2.** Shared annotated peaks within 3 kb of TSS.

**Supplementary Table 3.** Fusion-specific annotated peaks within 1 and 3 kb of TSS in bulk RNAseq

